# MuMu: a sample multiplexing protocol for droplet-based simultaneous single nuclei RNA- and ATAC-seq systems

**DOI:** 10.1101/2024.11.27.625728

**Authors:** Zhen Li, Tarik F. Haydar

## Abstract

Sample multiplexing is a common approach to reduce experimental cost and technical batch effect. Here, we present a protocol that for the first time allows the pooling of single nuclei from multiple biological samples prior to performing simultaneous single nuclei RNA-seq and ATAC-seq, which we term *Mu*ltiplexed *Mu*ltiome (MuMu). We describe steps for assembling the custom Tn5 transposome, performing the transposition reaction, nuclei pooling, sequencing library preparation, and sequencing data pre-processing. This protocol will greatly reduce the cost of sn-Multiome.

**Graphical abstract:** 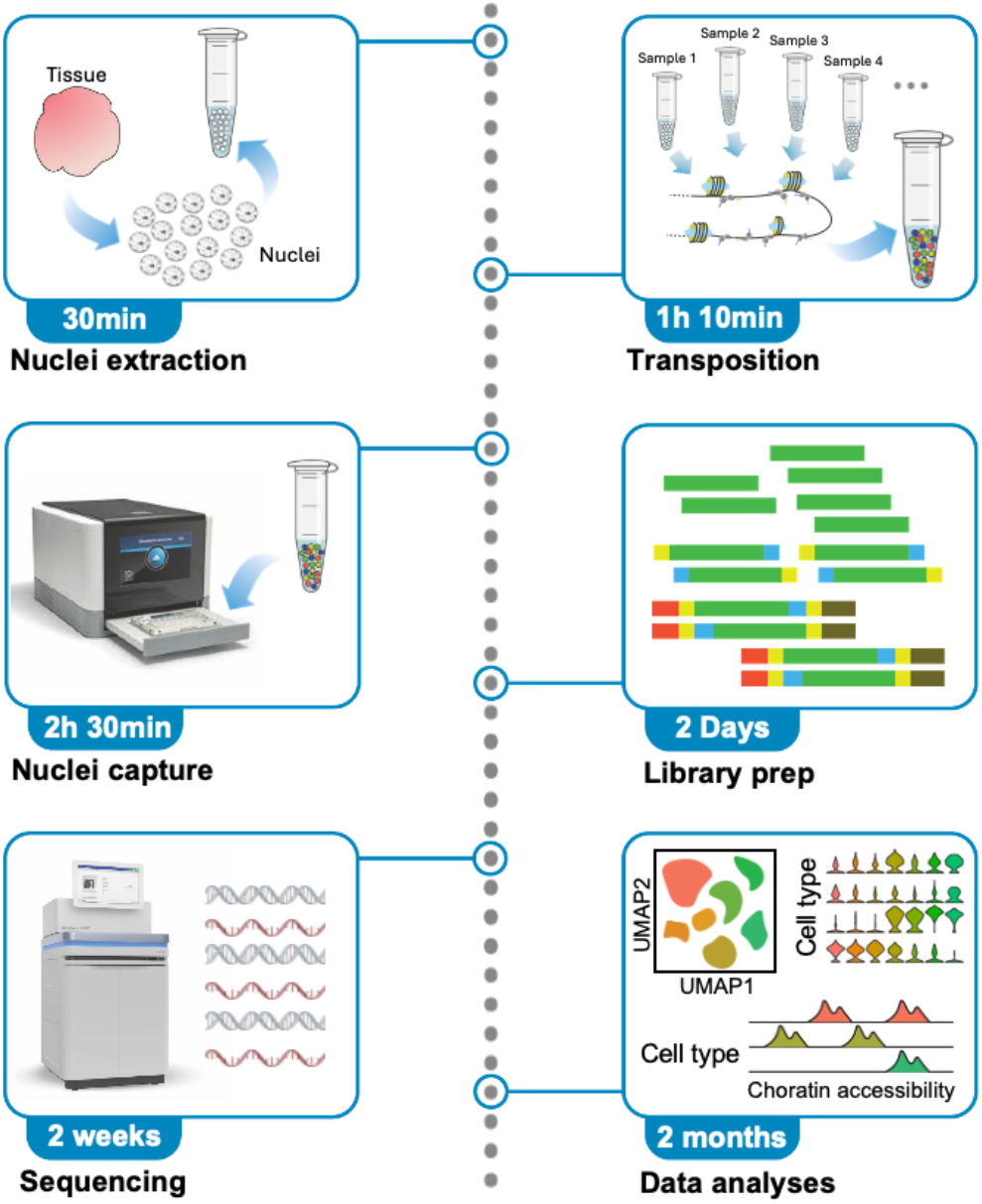

## Before you begin

The protocol below describes the specific steps for sample multiplexing before performing a sn-Multiome run with 10X Genomics Chromium Single Cell Gene Expression + ATAC kit. Sample multiplexing in this protocol refers to the action of combining single nuclei from multiple independent biological sources before performing droplet-based single nuclei capture and sequencing library preparation in a single reaction as if all the nuclei come from the same sample, significantly reducing the cost of analyzing multiple independent samples. Single cell multi-omics technologies have rapidly advanced in recent years.^1^ Simultaneous single nuclei RNA and ATAC sequencing technology (referred to hereafter as multiome sequencing technology), in particular, is well established and commercially available.^3^ However, the cost to perform one sn-Multiome reaction is very high. Moreover, biological experiments usually require multiple samples from different time points or conditions, in multiple replications. As a result, the cost to complete a whole experiment quickly rises to a prohibitive amount for many laboratories. Furthermore, when only a small number of nuclei is available as input, the potential output of a sn-Multiome reaction is often under-utilized, exacerbating cost-inefficiency. With the release of high throughput single cell capture platforms such as the Chromium X series (10X Genomics), whose capture capacity goes up to millions of cells or nuclei per capture, a reliable sample multiplexing strategy is now even more relevant and valuable. Thus, we attempt to improve the utility of droplet-based sn-Multiome by redesigning the Tn5 transposon sequence to allow the insertion of a sample specific barcode during the initial transposition reaction of the ATAC workflow and prior to single nuclei capture. Later on, the sample-specific barcode can be reliably detected in the fastq files containing sequencing reads and easily incorporated into read alignment steps for demultiplexing purposes. While the current protocol is written based on and specifically for the 10X Genomics Chromium Single Cell ATAC + GEX Multiome kit, it can be easily adapted to other droplet-based sn-Multiome strategies, such as single nucleus ATAC-seq and CUT&TAG with minor modifications.^2^ The same approach can also be used to multiplex bulk ATAC-seq samples.

Before you start, it is advisable that:

1. Institutional permission and oversight information for the animal study should be obtained. To demonstrate the outcome of the protocol, we collected samples from mouse neocortices at embryonic day (E) 16 and from pig neocortex at gestational week (GW) 10. The use of the animals was approved by the Institutional Animal Care and Use Committee (IACUC) at Children’s National Research Institute.
2. In most cases, the protocol begins with a piece of frozen tissue or cell suspension. We typically collect tissues in cryopreservation vials (2ml), snap-frozen in liquid nitrogen and kept in −80°C freezer, which we assume as the starting point of the protocol. This allows for the independent storage of multiple samples that can be combined at the start of the Multiome workflow. Other methods of sample storage are applicable and should minimally affect the outcome of the procedure, as long as the integrity of genomic DNA and messenger RNA (mRNA) is ensured.
3. The current protocol requires the in-house assembly of transposome, which starts with naked Tn5 transposases. We recommend purchasing the enzyme from a reliable commercial source, but experimenters may also opt to produce homemade Tn5 transposase, which has the benefit of reducing overall experimental cost even further. In either case, the activity and purity of transposase should be confirmed before proceeding.
4. The most significant change we have made to a standard sn-Multiome workflow is the replacement of transposon with MuMu barcode oligomer. The key resource table contains four examples of MuMu barcode oligomers but an infinite numbers of such oligomers can be designed to meet the multiplexing need of specific experiments. The structure of the 42bp oligomers is: GTGCTCTTCCGATCTXXXXXXXXAGATGTGTATAAGAGACAG. Substitute “XXXXXXXX” with a barcode of choice. The length of the barcode does not need to be 8-bp. However, we noticed that when a transposon is 60 bp or longer, it forms concatenation during transposition reaction (Figure 1). The concatenated transposon repeats interfere with sequencing library preparation and cannot be removed simply by size selection without losing genomic fragments all together.

**Figure 1.**
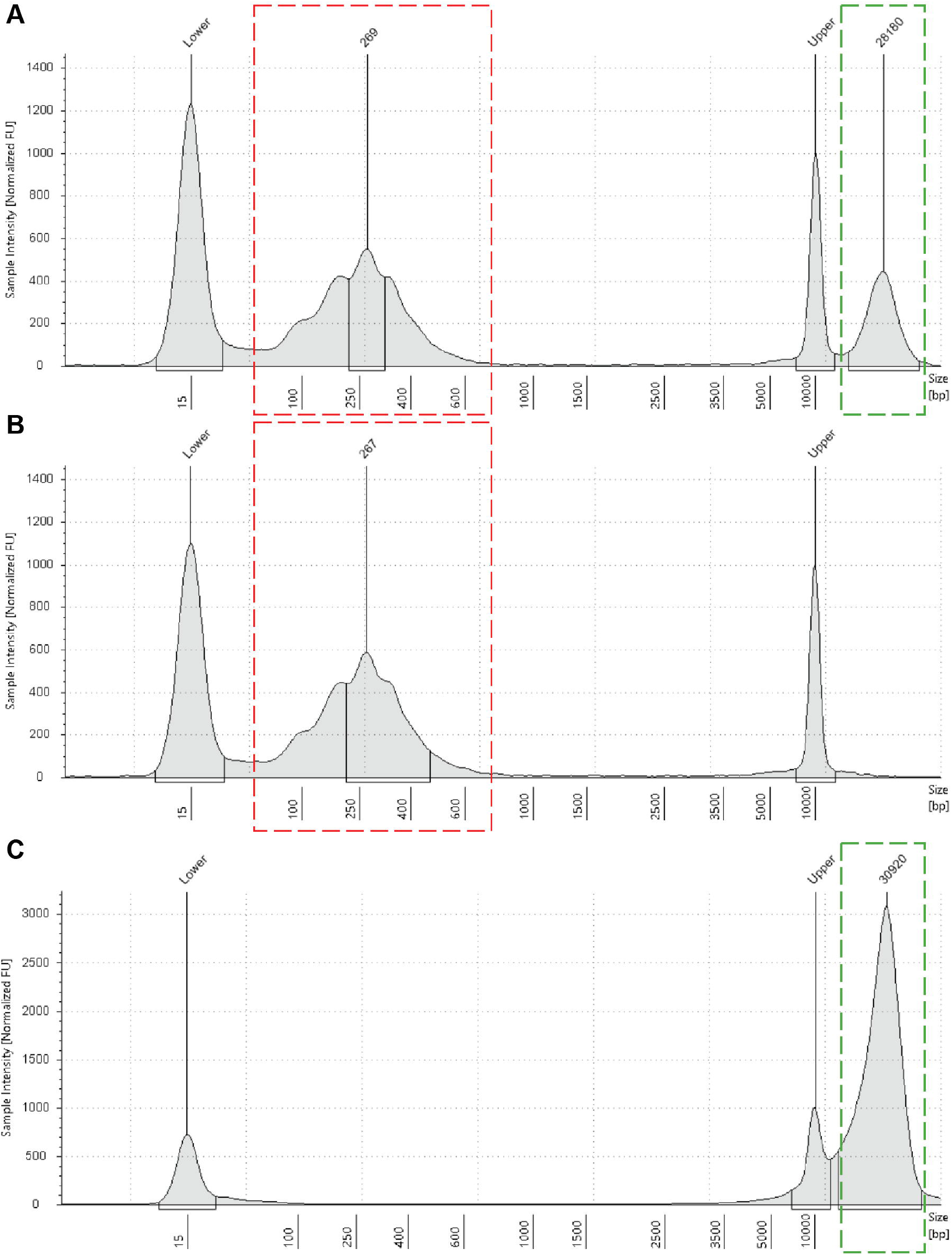
Tn5 transposome concatenate during reaction when transposon is 60 base pairs (bp) in size. (A and B) TapeStation analysis traces showing unexpected products (rectangle with red dashed line) with periodicity of roughly 60bp in the presence (A) or absence (B) of genomic DNA (rectangle with green dashed line), which the transposon is at 60bp. (C) No concatenation is observed using the MuMu barcode oligomers, which are 45 bp in size.

### Transposon preparation

#### Timing: 30min

1. Mix 5μL ME-A (100μM) with 5μL ME-Rev (100μM) and 40μL TE buffer in a PCR tube.
2. Mix 5μL of each MuMu barcoding oligomer (100μM) with 5μL ME-Rev (100μM) and 40μL TE buffer in PCR strips or plates. **Note:** The number of MuMu barcoding oligomers depends on the number of samples to be multiplexed. Prepare at least one barcoding oligomer for one sample. More than one barcoding oligomer can be used for one sample.
3. Put all solutions on thermocycler and run the following program:

| Step | Temperature | Time |
| --- | --- | --- |
| Denaturation | 98°C | 00:03:00 |
| Annealing | 70°C | 00:03:00 |
|  |  | Cooldown by 0.1 °C/sec |
| Hold | 25°C | forever |

### Transposome assembly

#### Timing: 35min

4 Combine the following reagents in a PCR tube and mix by pipetting:

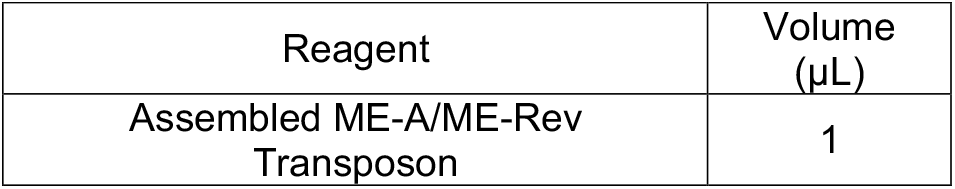
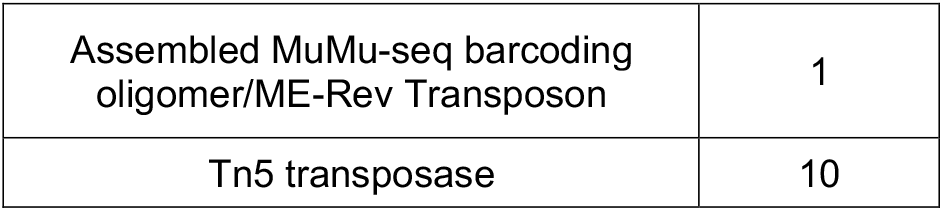 **Note:** The volumes used in this table are typically enough for four reactions, assuming around 10,000 input nuclei per reaction.
5 Incubate at 25°C in a thermocycler for 30min. **Note**: The assembled transposome can be kept at −20°C for at least four weeks.

### Prepare Pre-Amplification primers

#### Timing: 5min

6 Resuspend Pre-Amplification PCR primers to 100 µM.
7 Mix together 10µL of each primer and 60µL of TE buffer (100µL total).

### Key resources table

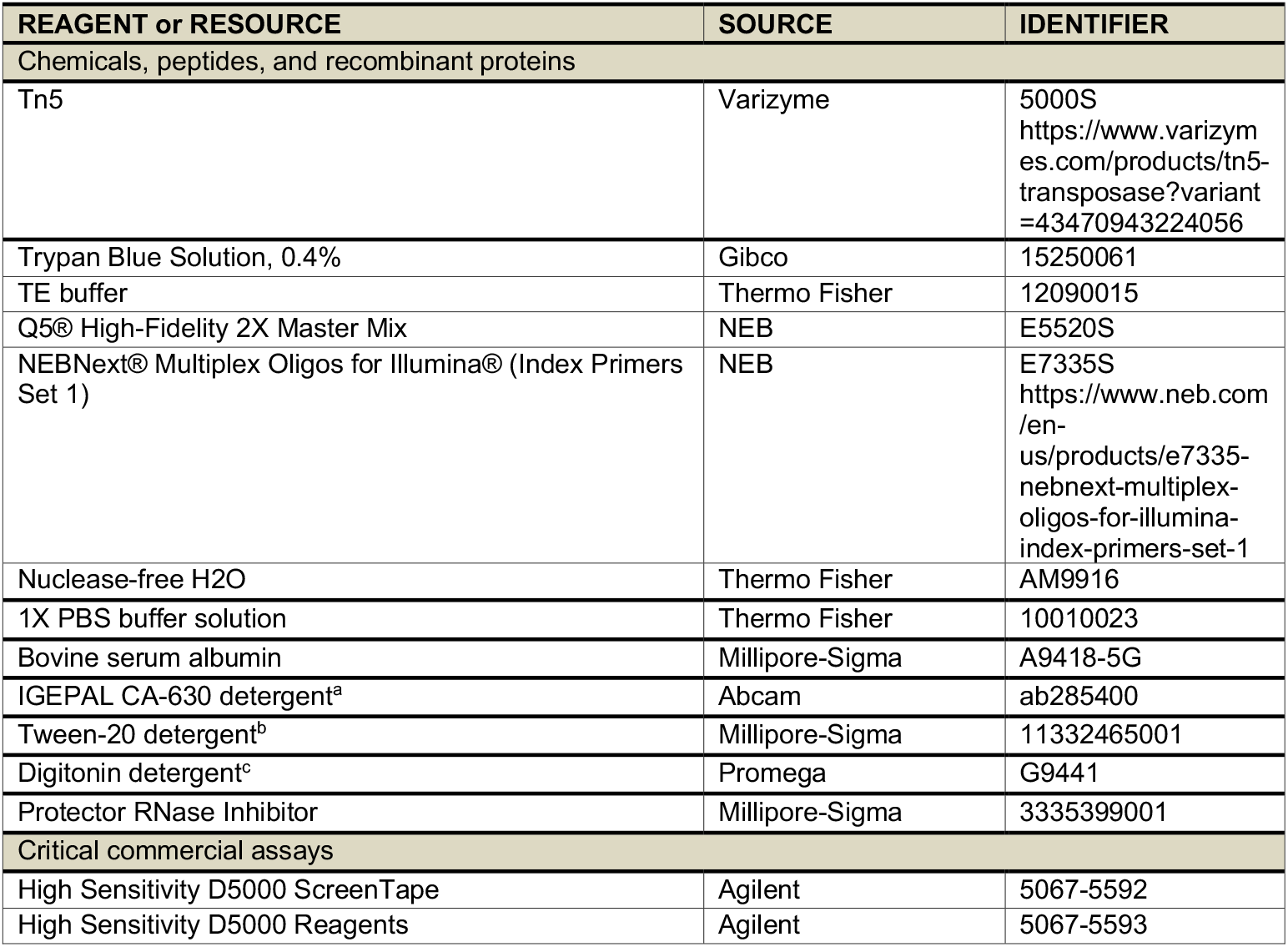
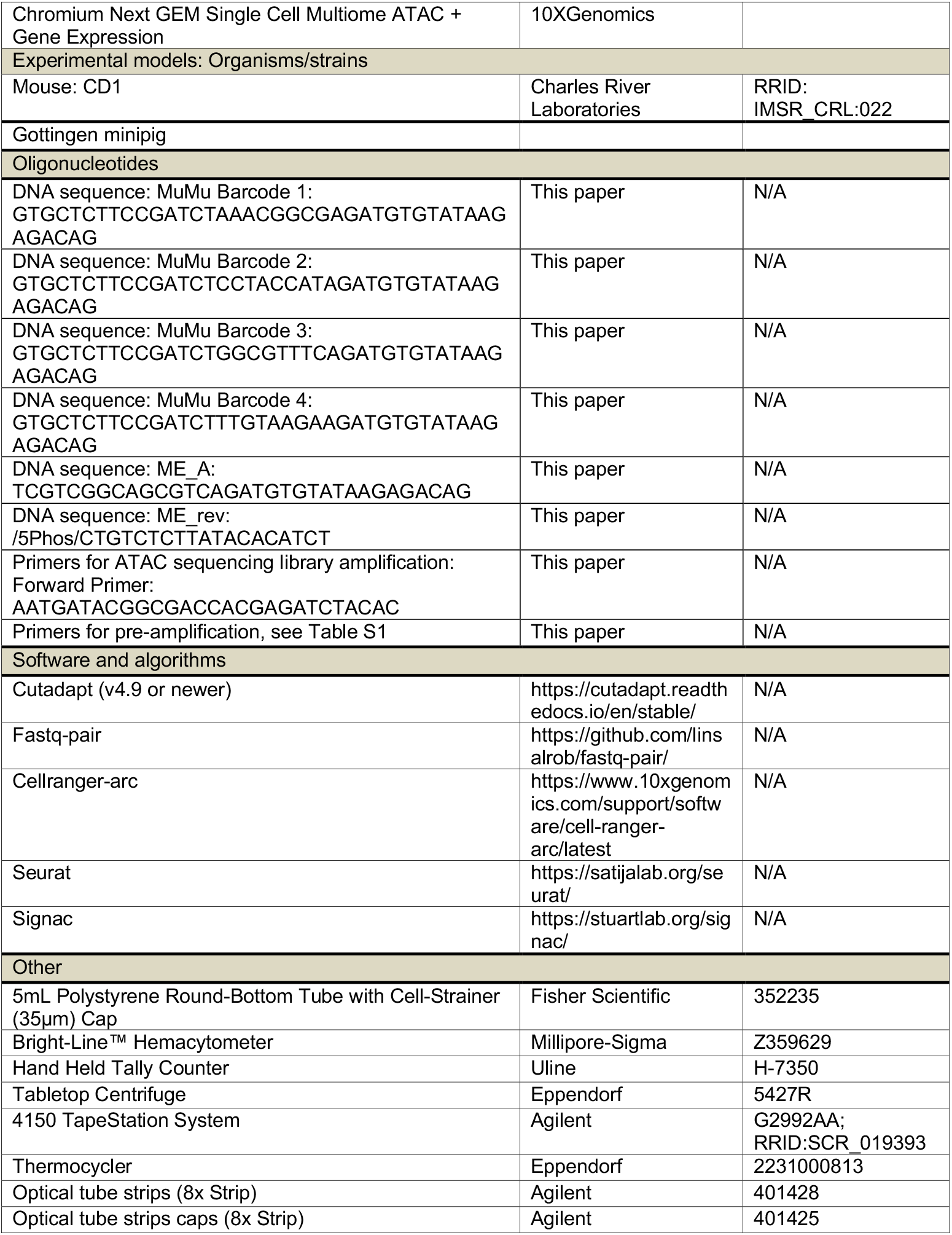
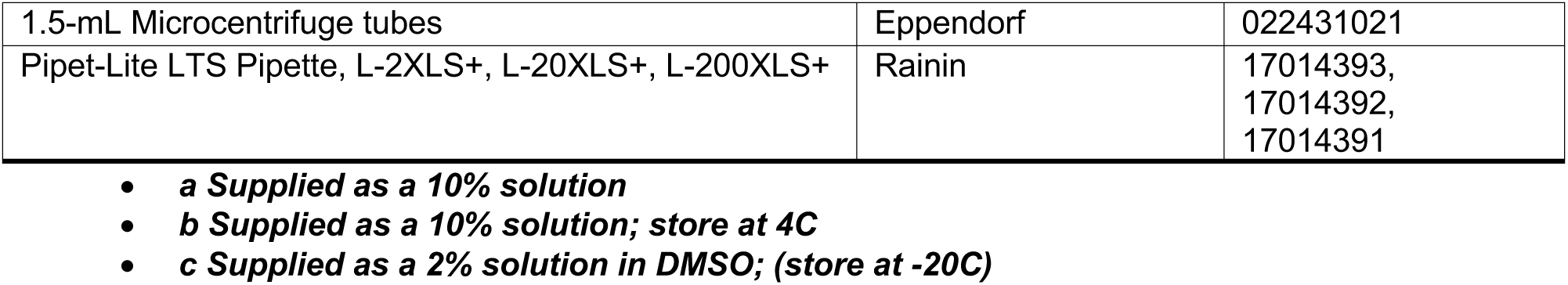

### Materials and equipment setup

#### Lysis buffer

| Reagent | Final concentration | Amount |
| --- | --- | --- |
| 1M Tris-HCL pH 7.4 | 10 mM | 500 µL |
| 5M NaCl | 10 mM | 100 µL |
| 1M MgCl<sub>2</sub> | 3 mM | 150 µL |
| 10% IGEPAL | 0.1% | 500 µL |
| 10% Tween-20 | 0.1% | 500 µL |
| 2% Digitonin | 0.01% | 250 µL |
| Protector RNase Inhibitor | 0.2 U/µL | 250 µL |
| 1X PBS buffer solution | n/a | 47.75 mL |
| **Total** | **n/a** | **50 mL** |

**Alternatives:** Use reagents provided by 10X Genomics Chromium Nuclei Isolation Kit.

##### Nuclei Resuspension Buffer

| Reagent | Final concentration | Amount |
| --- | --- | --- |
| Bovine serum albumin powder | 1% (wt/vol) | 0.5g |
| 0.5M Ethylenediaminetetraacetic acid (EDTA) | 1mM | 100 µL |
| Protector RNase Inhibitor | 0.2 U/µL | 250 µL |
| 1X PBS buffer solution | n/a | Up to 50 mL |
| **Total** | **n/a** | **50 mL** |

**Alternatives:** Use Nuclei Resuspension Buffer provided in the 10X Genomics Chromium Next GEM Single Cell ATAC + Gene Expression Multiome kit.

### Step-by-step method details

### Extract nuclei from each biological sample

#### Timing: 30min

This step extracts and isolates nuclei from frozen samples. The concentration of the final nuclei suspension is adjusted to a level that maximizes capture efficiency while minimizing the formation of doublets (capturing more than one nucleus per droplet).

1. Retrieve tissue or cells in a cryopreservation vial from −80°C freezer and immediately add 500µl of lysis buffer to the vial.
2. Incubate for 10min on ice.
3. After incubation, add 500µl nuclei resuspension buffer (RSB) and gently pipette 5-10 times until no visible cell lumps.
4. Adjust volume to 5ml and centrifuge at 800g for 5min to collect the nuclei.
5. Resuspend nuclei in RSB at appropriate volume (usually 1-2mL).
6. Filter resuspended nuclei into 5mL Polystyrene Round-Bottom Tube with Cell-Strainer Cap
7. Count nuclei with a hemacytometer.
  a. Mix 5µL nuclei suspension with 5µL Trypan Blue Solution 0.4% by gently pipetting at least 10 times.
  b. Load the mixture into the chamber on one side of the hemacytometer.
  c. Count the number of nuclei under a microscope and calculate the concentration.
8. Adjust nuclei concentration to about 1,000 nuclei/µL.

### Transposing genomic DNA in the extracted nuclei with MuMu barcode

#### Timing: 1h 10min

This step transposes the open chromatin regions of genome within each nucleus and simultaneously adds the sample specific barcode. At the end of this step, the nuclei from multiple barcoded samples are combined and processed in a single reaction. As an example, we assume an experiment with four biological samples.

9 For each sample, combine the following reagents in a 1.5 mL microcentrifuge tube separately:

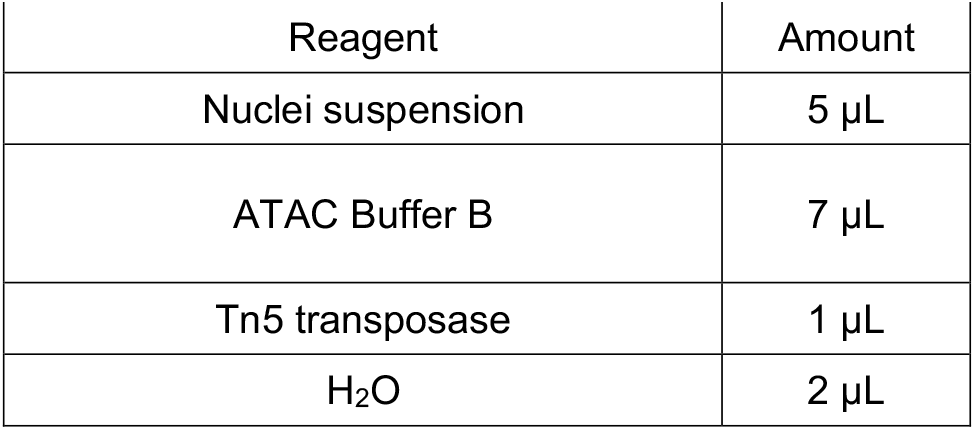 **Note:** If multiplexing more than four samples, adjust the number of input nuclei from each sample so that the final combined number does not exceed the maximum input amount required by the specific platform.
10 Incubate mixtures at 37°C for 1h on a thermocycler.
11 During incubation, prepare Master Mix.
  a. Combine reagents according to 10X Genomics Chromium Next GEM Single Cell Multiome ATAC + Gene Expression User Guide, Section 2.1a.
  b. Add 10 μL RSB to Master Mix and keep mixture on ice **Note:** If using 10X Genomics Chromium Next GEM Single Cell Multiome ATAC + Gene Expression kit, bring out Gel Bead from −80 °C freezer to equilibrate to room temperature.
12 Collect post-transposition nuclei by centrifugation.
  a. After the reaction, the nuclei in each tube are pelleted by centrifugation at 800g for 5 min. **Critical:** Record the tubes’ orientation in the centrifuge. A pellet may not be visible but nuclei will be collected on the outside wall of the tube after centrifugation.
  b. Using a P20 pipette, remove all supernatant by pipetting from the opposite side of the pellet.
  c. Gently wash the outside wall of each centrifuge tube using the master mix prepared in step 3 and combine sequentially.
  d. Keep master mix containing all the transposed nuclei on ice.

### Capture

#### Timing: 2h 30min

This step encapsules single nuclei within the droplet emulsions. Pre-mRNA and transposed genomic DNA fragments from each nucleus are released by lysis and captured on a gel bead. During reverse transcription, cell barcodes and unique molecular identifier (UMI) are added to mRNA (cDNA) and to genomic DNA fragments. Genomic DNA fragments and cDNA within the same droplet share the same cell barcode. Finally, the emulsion is broken and all molecules are collected for amplification in the next step.

13 The single cell capture is conducted faithfully following 10X Genomics Chromium Next GEM Single Cell Multiome ATAC + Gene Expression User Guide, Section 2.2 to Section 3.2.

**Pause point:** The product can be stored at 4 **°**C over night before proceeding to the next step.

### Pre-amplification

#### Timing: 35min

This step amplifies the genomic DNA fragments and cDNA from the previous step. Specifically for genomic DNA fragments, additional sequences are added at the end as primer binding sites for sequencing library amplification.

14. Perform PCR reaction as follows:

**PCR reaction master mix**

| Reagent | Amount |
| --- | --- |
| Product from Step 14 | 46 µL |
| Pre-amplification primers | 4 µL |
| Q5® High-Fidelity 2X Master Mix | 50 µL |

**Note:** Do not use any “hot start” polymerase since the PCR reaction begins with an initial extension step.

**PCR cycling conditions**

| Steps | Temperature | Time | Cycles |
| --- | --- | --- | --- |
| Initial Extension | 72 °C | 5 min | 1 |
| Initial Denaturation | 98 °C | 3 min | 1 |
| Denaturation | 98 °C | 20 sec | 7 cycles |
| Annealing | 58 °C | 30 sec |  |
| Extension | 72 °C | 1 min |  |
| Final extension | 72 °C | 1 min | 1 |
| Hold | 12 °C | forever |  |

15. Post pre-amplification cleanup: Follow 10X Genomics Chromium Next GEM Single Cell Multiome ATAC + Gene Expression User Guide, Section 4.3 to perform cleanup.

**Pause point**: Store at 4°C for up to 72 h or at −20°C for long-term storage.

### ATAC Sequencing Library Amplification

#### Timing: 20 min

This step amplifies the genomic DNA fragments and adds sequencing adaptors at the end.

16 Perform PCR reaction as follows:

**PCR reaction master mix**

| Reagent | Amount |
| --- | --- |
| Product from Step 15 | 40 µL |
| NEB Index | 5 µL |
| ATAC Forward Primer (10 µM) | 5 µL |
| Q5® High-Fidelity 2X Master Mix | 50 µL |

**PCR cycling conditions**

| Steps | Temperature | Time | Cycles |
| --- | --- | --- | --- |
| Initial Denaturation | 98 °C | 30 sec | 1 |
| Denaturation | 98 °C | 10 sec | 7 cycles |
| Annealing | 70 °C | 20 sec |  |
| Extension | 72 °C | 30 sec |  |
| Final extension | 72 °C | 1 min |  |
| Hold | 12 °C | forever | 1 |

### ATAC Sequencing Library Post-amplification Cleanup

#### Timing: 20 min

In this step, SPRISelect beads are used to purify fragments with a double-sided size selection strategy. The final library consists of fragments mostly from open chromatin regions of the genome, with a minor population of single and multi-nucleosome fragments.

17. Cleanup amplified ATAC sequencing library with SPRIselect beads.
  a. Vortex to resuspend SPRIselect reagent. Add 60 μL SPRIselect reagent (0.6X) to each sample. Pipette mix.
  b. Incubate 5 min at room temperature.
  c. Place on the magnet until the solution clears.
  d. Transfer 150 μL supernatant to a new strip tube. DO NOT discard the supernatant.
  e. Vortex to resuspend SPRIselect reagent.
  f. Add 65 μL SPRIselect reagent (1.25X) to each sample (supernatant). Pipette mix.
  g. Incubate 5 min at room temperature.
  h. Place on the magnet until the solution clears.
  i. Remove the supernatant.
  j. Add 300 μL 80% ethanol to the pellet. Wait 30 sec.
  k. Remove the ethanol.
  l. Add 200 μL 80% ethanol to the pellet. Wait 30 sec.
  m. Remove the ethanol.
  n. Centrifuge briefly. Place on the magnet.
  o. Remove remaining ethanol.
  p. a. Remove from the magnet. Immediately add 20.5 μl Buffer EB. Pipette mix.
  q. Incubate 2 min at room temperature.
  r. Centrifuge briefly. Place on the magnet until the solution clears.
  s. Transfer 20 μL sample to a new tube strip. **Pause point**: Store at 4°C for up to 72 h or at −20°C for long-term storage.

### Gene Expression Sequencing Library Preparation

#### Timing: 2 days

This step amplifies the cDNA library, which is subsequently fragmented. Then, the cDNA fragments are ligated with sequencing adaptors and amplified again through PCR reaction to generate the final sequencing library.

18 Follow 10X Genomics Chromium Next GEM Single Cell Multiome ATAC + Gene Expression User Guide, Section 6 and 7 to generate gene expression sequencing library.

### Sequencing Library Quality Control

#### Timing: 30 min

This is a routine quality control step before libraries are sequenced. Successful ATAC and gene expression sequencing libraries show typical size profiles of their corresponding library type. Refer to 10X Genomics Chromium Next GEM Single Cell Multiome ATAC + Gene Expression User Guide, Appendix: Agilent TapeStation Traces section for more details.

- We recommend using an Agilent TapeStation with the High Sensitivity D5000 kits to perform quality control on the final ATAC and gene expression sequencing libraries. Follow the manufacturer’s instructions depending on the exact instrument and kit used (TapeStation online protocol).

### Sequencing

#### Timing: 2-4 weeks

20 We recommend sequencing both gene expression and ATAC libraries with a paired-end, dual index protocol.

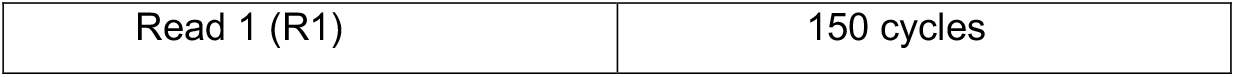
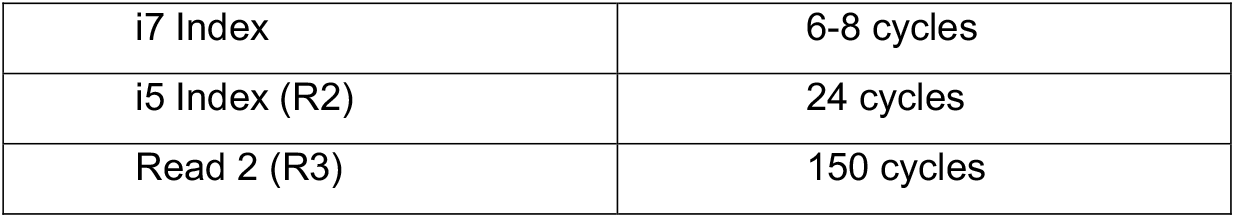 **Note:** The i7 index corresponds to the NEB index and is only relevant during the fastq file generation step. The NEB index is 6-bp in length. If 8-bp is required, append “AT” at the end.
21 Provide the NEB index sequence from step 16 to generate fastq files with bcl2fastq function.
22 Three fastq.gz files will be generated after sequencing. The file names should be adjusted according to Cell Ranger ARC requirement as follows(‘X’ indicates a single digit): Read 1: [Run name]_SX_LXXX_R1_XXX.fastq.gz I5 index: [Run name]_SX_LXXX_R2_XXX.fastq.gz Read 2: [Run name]_SX_LXXX_R3_XXX.fastq.gz

### Pre-process MuMu barcode sequences before read alignment

#### Timing: 2h

In this step, the fastq files are processed to prepare for alignment with the cellranger-arc function.

23. Generate sample specific fasta files corresponding to each MuMu barcode:

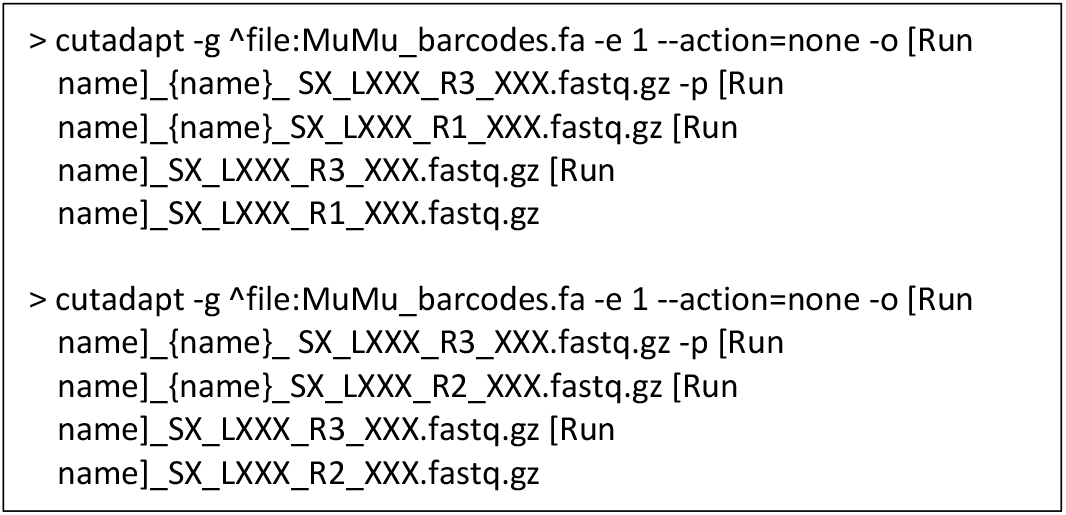 **Note:** {name} will be replaced by sample name supplied in the MuMu_barcodes.fa, which can be modified by the user.
24. Make sure you have three sets of ATAC fastq files for each of the MuMu barcoded samples: [Run name]_{name}_SX_LXXX_R1_XXX.fastq.gz [Run name]_{name}_SX_LXXX_R2_XXX.fastq.gz [Run name]_{name}_SX_LXXX_R3_XXX.fastq.gz
25. Run “cellranger-arc count” program for each sample, using the sample specific ATAC fastq files and combined gene expression fastq files as input: cellranger-arc count \ --reference={genome_reference} \ --id=[Run name]_{name} \ --libraries=libraries.csv
26. (Optional) Run cellranger-arc aggr to aggregate and normalize all samples into one dataset.

### Expected outcomes

The sn-Multiome ATAC library generated with the MuMu protocol is identical to ones that are generated with the standard 10X Genomics Chromium Next GEM Single Cell Multiome ATAC + Gene Expression workflow (Figure 2). Sometimes a smaller peak (less than 150 bp) appears in the final library, which represents adaptor dimer and should not affect sequencing or downstream analyses.

**Figure 2.**
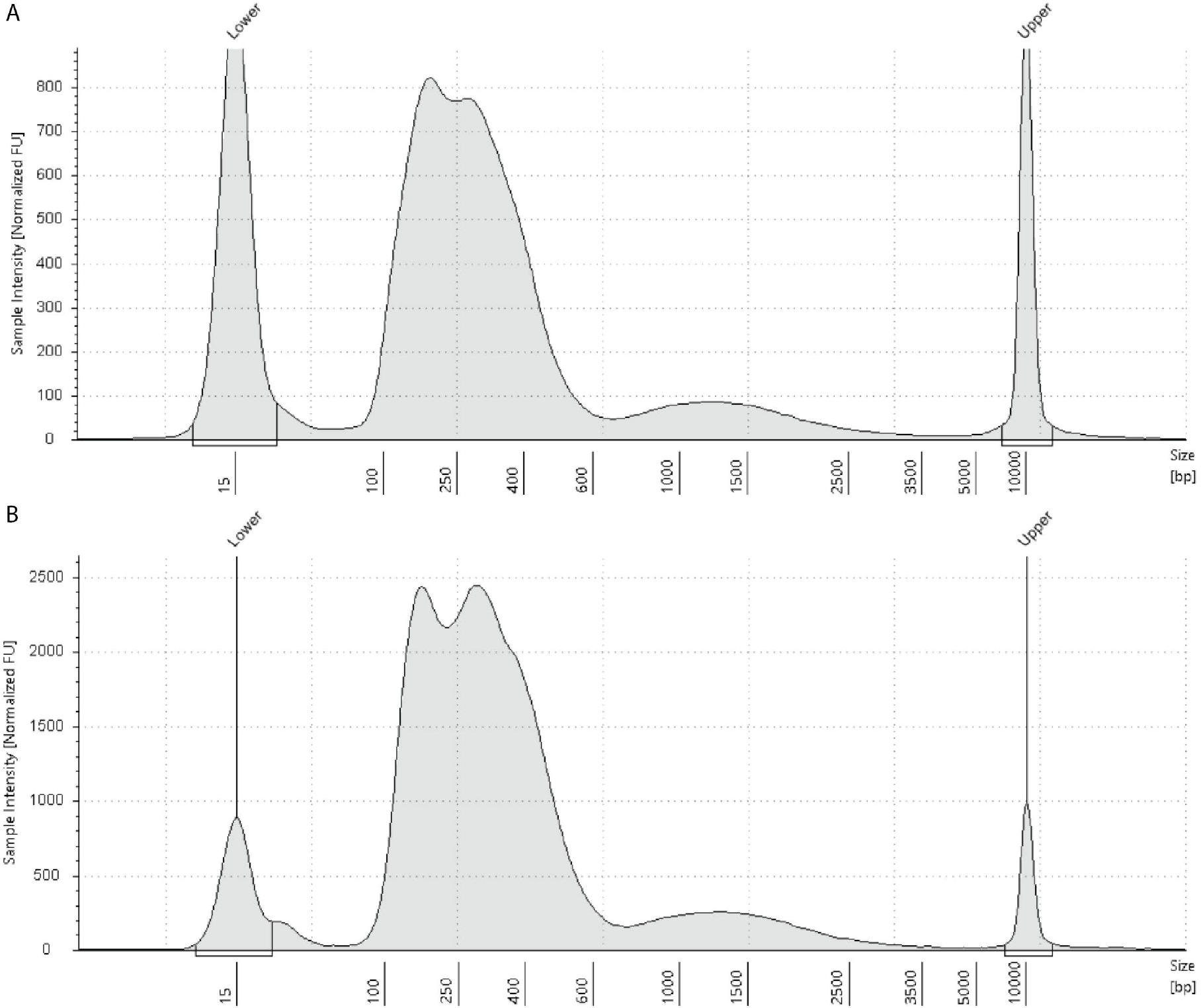
Typical TapeStation traces of finished ATAC libraries produced using (A) the current protocol or (B) the original 10X Genomics Chromium Next GEM Single Cell Multiome ATAC + Gene Expression protocol.

### Quantification and statistical analysis

As a demonstration of the MuMu protocol, we processed tissue samples obtained from a mouse neocortex at E16 and a pig neocortex at gestation week (GW) 10. The nuclei from the two pieces of tissue samples were extracted. We then performed transposition reaction with the MuMu barcode oligomers as the transposon on two separate pools of nuclei from each tissue sample. The two mouse nuclei pools correspond to sample 1 and sample 3, while the pig nuclei pools correspond to sample 2 and sample 4, respectively. Following the MuMu preparation and bioinformatics preprocessing pipeline, more than 90% of all the raw ATAC reads contained correct MuMu barcodes and were confidently assigned to their corresponding sample (Figure 3A). We first assessed the rate at which MuMu barcodes mix with one another. When RNA or ATAC reads from any nucleus are mapped to the appropriate genome reference (mouse-to-mouse or pig-to-pig), the alignment rate is around 90%. However, when reads are mapped to the opposing reference, the alignment rate is only around 20%, due to heterology between mouse and pig genomes. We demonstrated this pattern by mapping a mouse sn-Multiome dataset (“10X Multiome”) generated in-house using the original 10X Genomics Multiome protocol to both the mouse and the pig genome references (Figure 3B).

**Figure 3.**
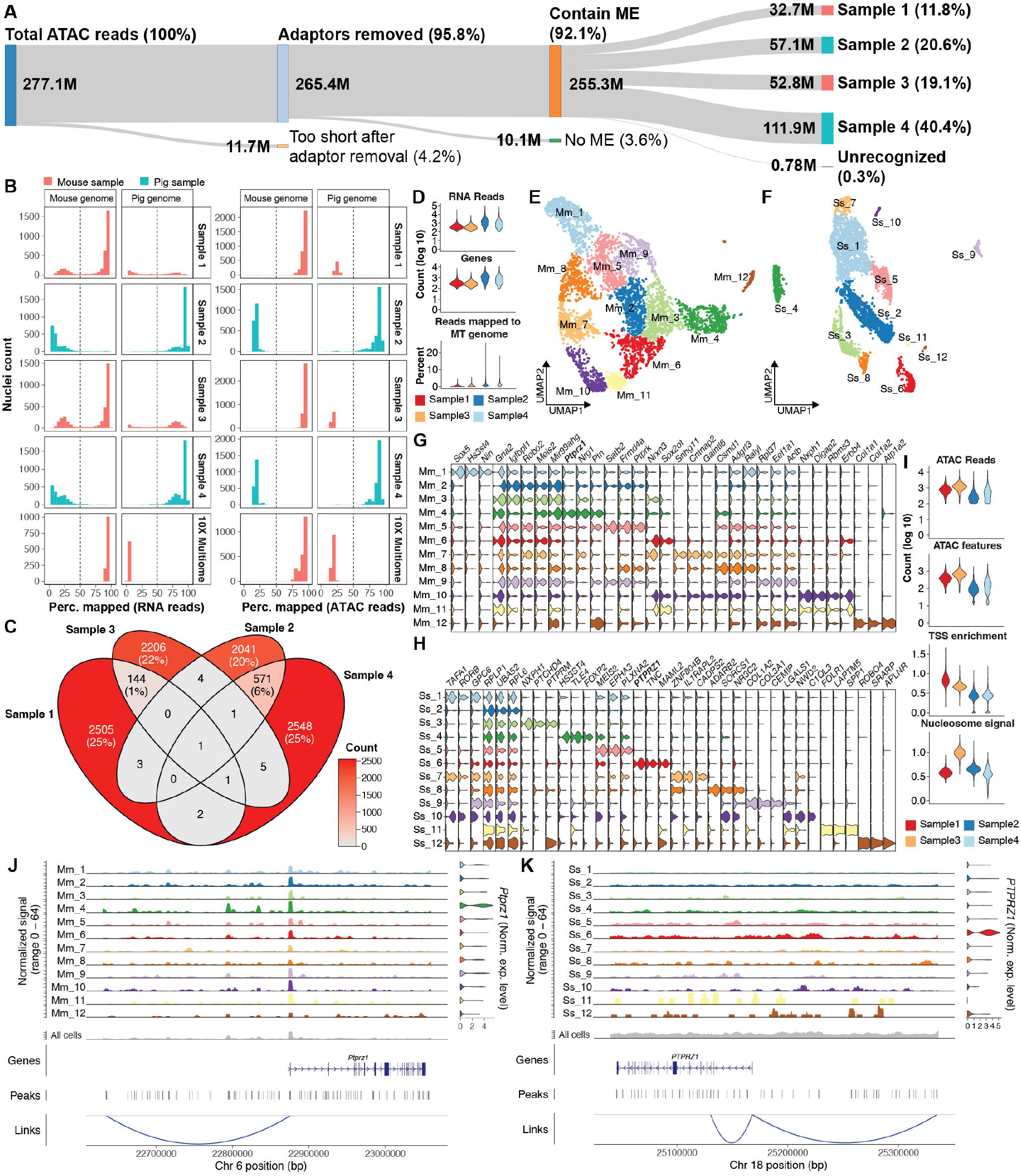
Demonstration of sample multiplexing and downstream data analyses. (A) Sankey diagram showing the breakdown of all ATAC reads acquired through sequencing at each data preprocessing step. The percentage out of the total ATAC reads for each category is labeled in parentheses. ME, mosaic end; M, million reads. (B) Histograms showing the distribution of percentages of RNA or ATAC reads mapped to mouse or pig genome from each nucleus. Sample 1 and 3 are collected from a mouse neocortex at embryonic day (E) 16, while 2 and 4 are from a pig neocortex at gestation week 10. Dotted vertical lines indicate 50% mapping rate (“cutoff”). Additionally, an E15.5 mouse neocortex sample (“10X Multiome”) processed using the original 10X Genomics Chromium Next GEM Single Cell Multiome ATAC + Gene Expression kit was included for comparison. (C) Venn diagram showing number of cell barcodes presented in each sample. Cell barcodes presented in more than one sample are likely “doublets” and are removed from downstream analyses. (D) Violin plots showing distributions of RNA read counts, detected genes, and percentage of reads mapped to mitochondrial genome in nuclei passing quality control. MT, mitochondria. (E and F) Uniform manifold approximation and projection constructed with gene expression data showing single nuclei profiles from the mouse (C) or the pig (D) samples. Colors represent cell clusters identified through the Seurat analysis pipeline. (G and H) Violin plots showing the most specifically expressed genes associated with each cell cluster of the mouse (E) or the pig (F) samples. Clusters are arranged by size from largest to smallest. (I) Violin plots showing distributions of ATAC read counts, ATAC features, transcription start site enrichment, and nucleosome signal. TSS, transcription start site. (J and K) Chromatin accessibility profiles around gene *PTPRZ1* in each cell cluster of the mouse (J) or the pig (K) samples. *PTPRZ1* expression levels are shown on the right as violin plots. The accumulative chromatin accessibility profile from all nuclei is shown below in dark grey., followed by a diagram of *PTPRZ1* gene body. All identified ATAC peaks (“Peaks”) within the region are shown as vertical lines. A peak in the upstream region (“Links”) is identified in both mouse and pig genome that is correlated with gene expression of *PTPRZ1*. Norm. exp. level, normalized expression level.

Based on this observation, we reasoned that if there is frequent mixing between MuMu barcodes, one would expect a substantial shift in the distribution of mapping ratio and that more nuclei would be assigned to one species but have their reads mapped to the genome of the opposing species. Our analysis indicated that all the nuclei displayed the correct pattern of mapping ratio based on the percentage of ATAC reads mapped to the two genomes, suggesting that there is no mixing or interchange of the MuMu barcodes after the transposition reactions. While RNA reads from a small number (4.3%) of the total nuclei mapped mainly to the opposing genome, none of these nuclei had a mapping rate of the ATAC reads to the opposing genome at more than 50%, an average of 20.1% (SE±0.1%). The erroneously mapped RNA reads were most likely due to ambient RNA. This phenomenon can be used to exclude this minority of nuclei. Together, our data demonstrated that the MuMu barcodes faithfully and specifically labeled the samples they were assigned to. The nuclei with more than 50% of either RNA or ATAC reads mapped to the opposing genome were removed from downstream analyses.

A unique advantage of the MuMu barcoding system is its ability to unambiguously detect doublets, due to its reliability. Thus, we next examined how many cell barcodes (CB) were assigned to more than one sample as an indication of doublets. We found 144 CB shared between Sample 1 and Sample 3, as well as 571 CB shared between Sample 2 and Sample 4 (Figure 3C). The doublet rates were therefore about 5% between the mouse samples and about 20% between the pig samples. While the doublet ratio of the pig samples was higher than the mouse samples, it was still within the expected range of a droplet based single cell experiment^4^, and may reflect an underestimation of the nuclei concentration during the extraction procedures for one of the pig samples (Sample 4) due to human error. We removed any CB that was assigned to more than one sample from downstream analyses.

The remaining nuclei displayed compatible distributions of RNA read counts, detected gene counts and percentage of RNA reads mapped to mitochondrial genome to standard single cell RNA-seq datasets. We then performed the Seurat (v5) analyses pipeline^5^ and identified cell clusters in uniform manifold approximation and projection graphs (Figure 3E and F). The cell clusters from either species expressed the expected marker genes appropriate for the neocortical developmental period. Notably, E16 in mouse and GW10 in pig mark the beginning gliogenesis in the two species (Figure 3G and H). As expected, we identified oligodendrocyte precursor cell populations (Mm_2 and Ss_8) in both datasets, which were marked by a specific expression of *PTPRZ1*. The ATAC data also displayed compatible quality to standard single cell ATAC-seq datasets, reflected by the distributions of ATAC read count, number of ATAC features, transcription start site enrichment and nucleosome signal. We then examined the chromatin accessibility landscape around the gene body of *PTPRZ1*, we observed an enhanced openness of the *PTPRZ1* locus. More importantly, in both species, we were able to identify an ATAC peak about 200,000bp upstream of the gene that correlated with the *PTPRZ1* RNA expression (Figure 3J and K).^6^ The analyses thus identify a potentially common regulatory element of *PTPRZ1* shared by the two species.

### Limitations

The limitations commonly associated with snATAC-seq and snRNA-seq also apply to the protocol described here. We want to highlight one particular limitation that pertains to the transposition reaction. Current state-of-the-art ATAC-seq protocols, including the one described here, utilize transposon oligomers with two different sequences (mixed at equal molarity), so that DNA fragments can be tagged with different oligomers on each end to serve as primer binding sites for the PCR amplification. As a result, however, about half of the transposition events generate DNA fragments with the same oligomer sequences at both ends, and these “homo-tagged” fragments cannot be amplified. Thus, 50% of the transposed DNA fragments are lost by default.^7^ This is not a significant drawback in bulk-tissue ATAC-seq because there are presumably many copies of the genome from the large amount of nuclei with the same open chromatin configuration. However, at the single nucleus level, each genome is considered unique, which means the open chromatin fragments from each nucleus are also unique. The loss of 50% fragment from each single nucleus is irreversible and poses a challenge on the interpretation of final sequencing data.

A limitation specific to the current protocol is the randomness in the sequencing read distribution and nuclei distribution among the MuMu barcoded samples, which is affected by many factors. The sequencing and nuclei distributions are only accessible after sequencing, at which point it is already too late to make an adjustment. This limitation emphasizes the importance of accurate estimation of nuclei concentration during the extraction steps and precise techniques in pipetting nuclei between steps. The transposition reaction, especially the amount of transposome used, should also be carefully tested to achieve the most optimal number of fragments from open chromatin regions. It may be advisable to always perform a test sequencing run at low read depth to see the distributions before committing to a high throughput run.

## Troubleshooting

### Problem 1

Failed to see a multimodal ATAC sequencing library profile. Only one peak can be detected at around 200 bp (Step 19).

This type of library profile usually indicates degradation of chromatin structure, resulting from poor quality of starting material. It is also possible that too much transposome is used. Another possible cause is an error during ATAC sequencing library post-amplification cleanup.

### Potential solution

- Collect fresh tissue. Be sure to snap-frozen sample as soon as possible in liquid nitrogen and transfer it to −80 °C right away after freezing for long term storage.
- Make sure to perform all nuclei extraction procedure on ice. Centrifuge nuclei at 4 °C until transposition step.
- When in doubt, perform a bulk ATAC-seq with sample in question to assess quality before proceeding to sn-Multiome.
- Try reducing the amount of transposome during the transposition step.
- During ATAC sequencing library post-amplification cleanup, make sure SPRIselect beads are well mixed every time and use the recommended ratio of beads to sample.

### Problem 2

Nuclei clump after extraction and cannot be dissociated further (Step 1-8).

This is usually caused by over-lysis resulting in the release of genomic DNA from broken nuclei, due to excessive pipetting or strong lysis buffer.

### Potential solution

- Pipette gently during nuclei extraction steps and reduce pipetting frequency depending on tissue type.
- Reduce the amount of detergent in the lysis buffer.
- Add DNase I to lysis buffer may help digest free-floating genomic DNA and reduce tangling.

### Problem 3

The final concentration of ATAC sequencing library is low (Step 14-16). This usually indicates an overestimation of the number of captured nuclei.

### Potential solution

- Quantification of starting number of nuclei: The initial estimation of nuclei concentration needs to be accurate. Carefully count and calculate the concentration of nuclei after extraction.
- Sufficiently resuspend nuclei containing solution to homogeneity before mixing transposition reaction.
- Carefully collect nuclei after transposition with master mix.
- Further amplify sequencing library with Illumina P5 and P7 primers.

## Resource availability

### Lead contact

Further information and requests for resources and reagents should be directed to and will be fulfilled by the lead contact, Tarik F. Haydar, PhD

### Technical contact

Technical questions on executing this protocol should be directed to and will be answered by the technical contact, Zhen Li, PhD

### Data and code availability

The datasets described in this protocol can be found on SRA with accession number PRJNA1191389.

## Acknowledgments

This work was supported by NIH grants 1R01NS116418-01 and 5R01NS116418-02. The authors wish to thank the Genomics Core at the Center for Genomic Medicine, Children’s National Research Institute for providing access to equipment.

## Author contributions

Z. L. designed and executed all experiments, including bioinformatics analyses. T.F.H. supervised the project. Z. L. and T. F. H. composed the manuscript.

---

**Table 1:**
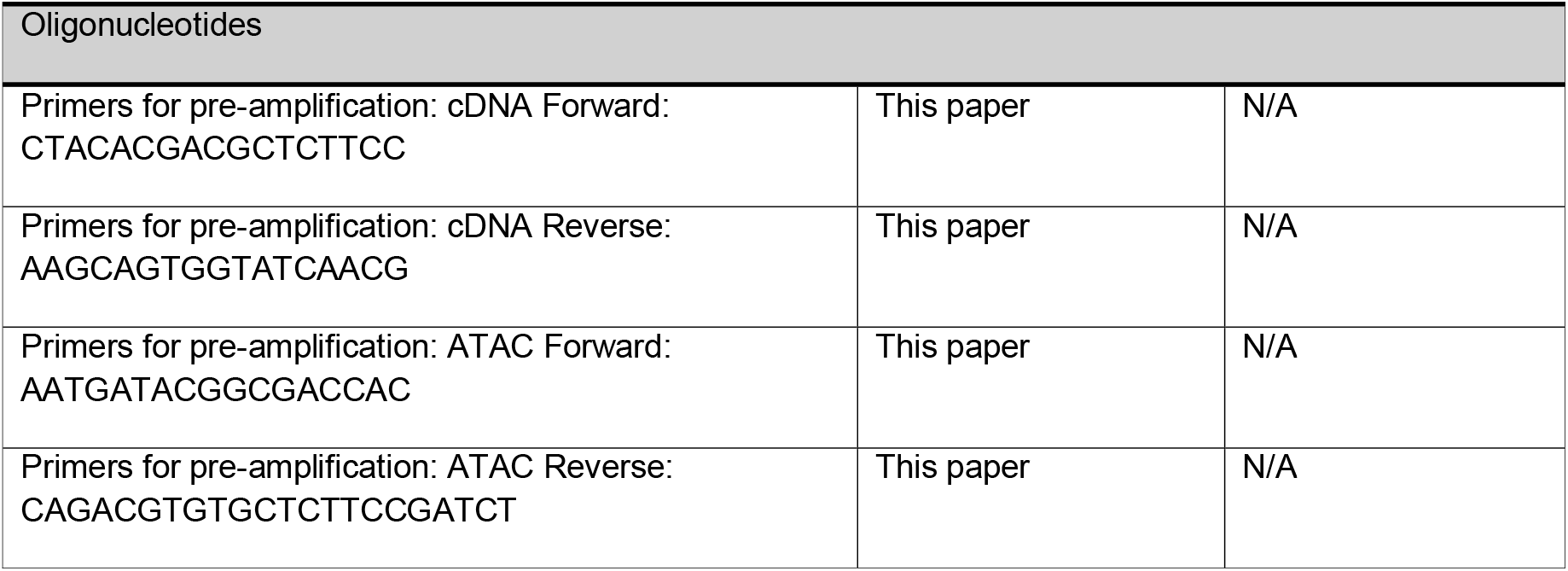
Pre-amplification primers.

